# Axonal localization of the Fragile X family of RNA binding proteins is conserved across mammals

**DOI:** 10.1101/464552

**Authors:** Katherine A. Shepard, Lulu I T. Korsak, Michael R. Akins

**Author notes:** **Correspondence to:** Michael R. Akins, Ph.D., Department of Biology, Drexel University, PISB 319; 3245 Chestnut St., Philadelphia, PA 19104. Current Address: Department of Neurology, University of Pennsylvania Perelman School of Medicine, Philadelphia, PA 19104 USA.

## Abstract

Spatial segregation of proteins to neuronal axons arises in part from local translation of select mRNAs that are first transported into axons in ribonucleoprotein particles (RNPs), complexes containing mRNAs and RNA binding proteins. Understanding the importance of local translation for a particular circuit requires not only identifying axonal RNPs and their mRNA cargoes, but also whether these RNPs are broadly conserved or restricted to only a few species. Fragile X granules (FXGs) are axonal RNPs containing the Fragile X related family of RNA binding proteins along with ribosomes and specific mRNAs. FXGs were previously identified in mouse, rat, and human brains in a conserved subset of neuronal circuits but with species-dependent developmental profiles. Here we asked whether FXGs are a broadly conserved feature of the mammalian brain and sought to better understand the species-dependent developmental expression pattern. We found FXGs in a conserved subset of neurons and circuits in the brains of every examined species that together include mammalian taxa separated by up to 160 million years of divergent evolution. A developmental analysis of rodents revealed that FXG expression in frontal cortex and olfactory bulb followed consistent patterns in all species examined. In contrast, FXGs in hippocampal mossy fibers showed an increase in abundance across development for most species except for guinea pigs and members of the *Mus* genus, animals that navigate particularly small home ranges in the wild. The widespread conservation of FXGs suggests that axonal translation is an ancient, conserved mechanism for regulating the proteome of mammalian axons.

## Introduction

Local protein synthesis is a broadly conserved mechanism for controlling spatiotemporal patterns of protein expression in both eukaryotic and prokaryotic cells and is critical for cell division, migration, and process elaboration (Martin and Ephrussi, 2009; Nevo-Dinur et al., 2011; Buxbaum et al., 2015). Restricting local translation to the appropriate cellular compartment at the correct developmental timepoint requires correct positioning of ribonucleoprotein particles (RNPs), complexes that contain mRNAs and the RNA binding proteins that control their translation. In neurons, local translation is supported by a variety of RNPs that can differ in their prevalence, mRNA cargoes, and RNA binding protein composition depending upon developmental stage, neuronal cell type, and subcellular location. The large number of neuronal RNPs reflects the diversity of neuronal morphology and function seen throughout the nervous system. However, since brain size and structure can vary drastically across species, some of these RNPs may support specialized functions in only one or a few species. The most interesting RNPs therefore are those whose expression is conserved across different species, since they are compelling candidates for supporting fundamental neurobiological functions.

Fragile X granules (FXGs) are a class of neuronal RNPs first identified in mice and rats that exhibit remarkable specificity in the circuits in which they are found, the RNA binding proteins they contain, and the mRNAs with which they associate (Christie et al., 2009; Akins et al., 2012, 2017; Chyung et al., 2018). Within the subset of neurons that express them, FXGs are found only in axons and are excluded from the somatodendritic domain (Christie et al., 2009; Akins et al., 2012, 2017). FXGs contain one or more of the Fragile X related (FXR) family of RNA binding proteins: FMRP (Fragile X mental retardation protein), FXR2P, and FXR1P. FMRP, which is mutated in the autism-related disorder Fragile X syndrome (FXS), modulates activity-dependent synaptic protein synthesis and therefore is a critical regulator of experience-dependent circuit refinement in both the developing and adult brain (Bassell and Warren, 2008; Darnell and Klann, 2013). Notably, FXGs associate with a specific subset of FMRP target mRNAs whose protein products modulate the structure and function of presynaptic sites (Akins et al., 2017; Chyung et al., 2018). These FXG properties are conserved in both mice and rats, suggesting these granules may support conserved cellular functions. FXG-regulated local protein synthesis may thus provide a conserved mechanism for regulating neuronal circuit function.

Interestingly, however, the developmental timing of FXG expression differs among species (Akins et al., 2017). In mice, FXGs are found in the olfactory bulb throughout life but in other forebrain regions are only found in juvenile animals during periods of robust experiencedependent circuit refinement. In contrast, FXGs are found in rats throughout life in both hippocampal and olfactory bulb circuits while recapitulating the developmental downregulation in other forebrain regions seen in mice. Importantly, FXGs are also present in adult human hippocampus in the same neuron types in which they are found in mice and rats. FXGs are thus positioned to regulate neuronal circuits during development as well as in adulthood.

To elucidate whether FXGs play fundamental roles in the mammalian brain, we asked whether they are conserved in a broad variety of mammalian species and, if so, whether their expression patterns are also conserved or instead vary among species. In particular, we sought to better understand the species-dependent expression of FXGs in the adult hippocampus. To these ends, we examined brain tissue from mammalian species that together cover 160 million years of divergent evolution (Fig. 1). FXGs were present in all these species in a spatial pattern equivalent to that seen in mice. Interestingly, most of the species expressed FXGs in the adult hippocampus. To better understand the developmental profile of FXG expression, we examined olfactory bulb, cortex, and hippocampus from juveniles and adults of five rodent species. Olfactory bulb and cortex had similar developmental patterns in all these species, even though the absolute abundance of FXGs was species-dependent. In contrast, FXGs were found in both juvenile and adult hippocampal circuits in voles and deer mice (*Peromyscus*) but were only abundant in juvenile hippocampus in all examined representatives of the *Mus* genus as well as in guinea pigs. Our results suggest that FXG-regulated local translation is a broadly conserved mechanism for regulating the axonal proteome of select neurons in the mammalian brain. Moreover, this regulation occurs in most species during both developmental and adult periods of experience-dependent circuit refinement.

**Figure 1.**
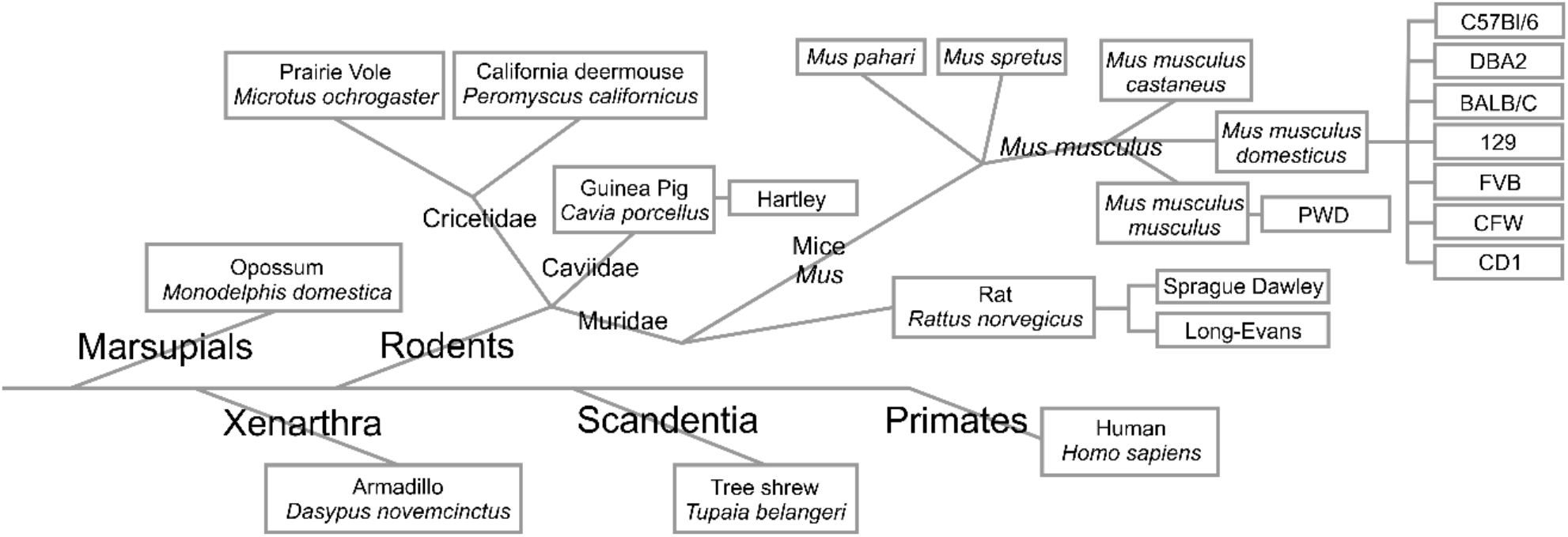
Taxonomic key of examined species. This key depicts the taxonomic relationships among the species investigated in the present study as well as in a past study (Akins et al., 2017).

## Results

### FXGs are highly conserved across mammalian taxa

We first asked whether FXGs are broadly conserved by examining hippocampal tissue from adults of a wide range of mammalian species. In the hippocampus of mice, rats, and humans, FXGs are found in two populations of axons – dentate mossy fibers and CA3 associational fibers – but absent from others including the Schaffer collaterals. To test the extent of FXG conservation, we immunostained hippocampal sections from adult animals for FXR2P, a necessary component of all FXGs (Christie et al., 2009), and FMRP. Consistent with past studies, FXGs were readily observed in mossy fibers in adults of most examined species, including tree shrews, armadillos, opossums, and voles (Fig. 2). We also observed FXGs in associational fibers in all of these species except opossum. Since we do not have an independent marker of these axons, we could not determine whether this axon population does not contain FXGs in opossums or whether the axons were not present in our specimens. These findings reveal an ancient origin for FXG expression as they were found in brains from a wide taxonomic range, including both placental and non-placental mammals.

**Figure 2.**
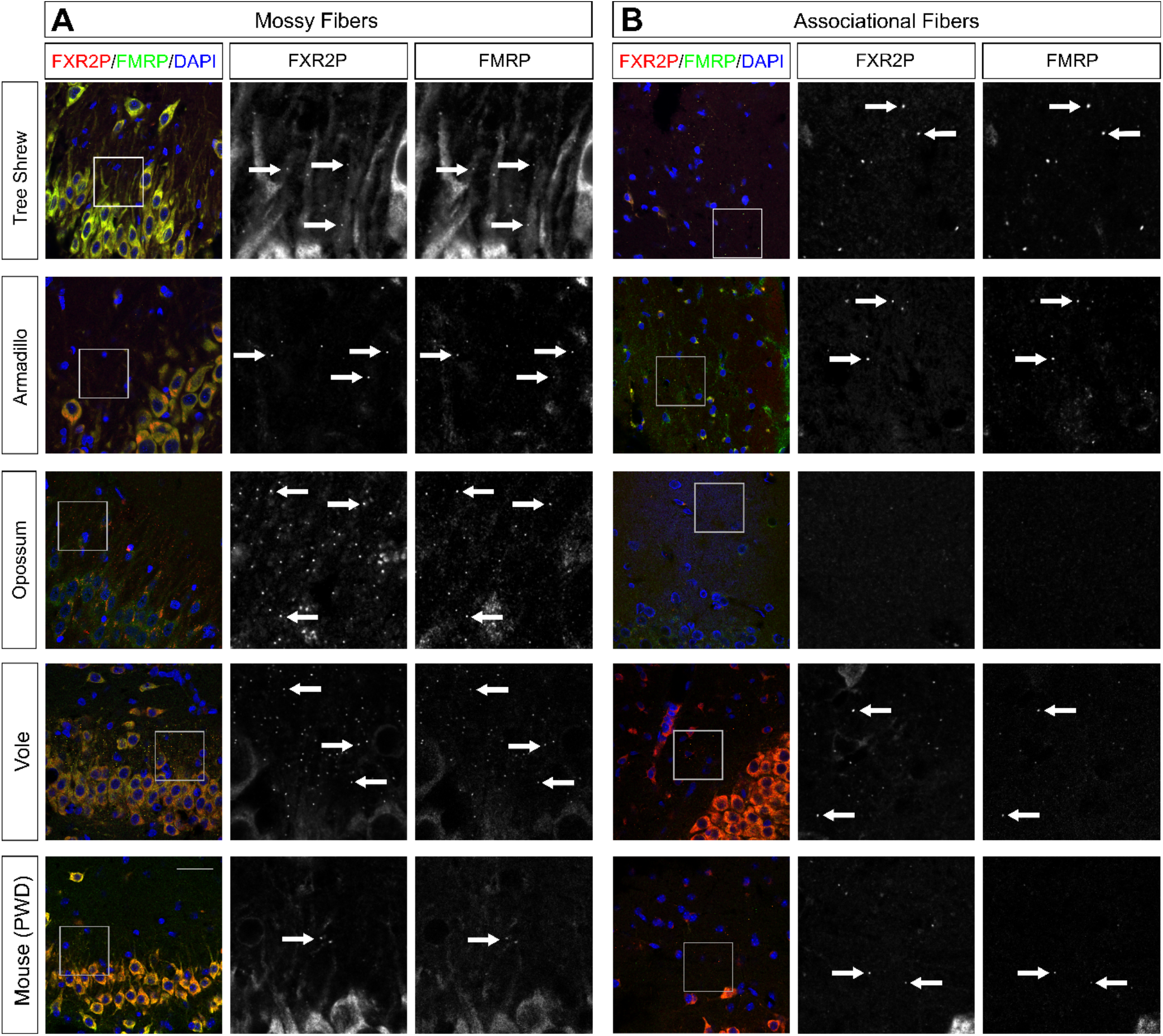
FXGs containing FMRP are expressed in the hippocampus in adults of divergent species. Confocal micrographs of immunostaining for FXR2P (red) and FMRP (green) in **(A)** mossy fibers and **(B)** associational fibers in hippocampal tissue from adult armadillo (exact age unknown), tree shrew (P322), opossum (P97), vole (P120), and mouse (PWD strain, P120). FXGs were identified in mossy fibers in all species examined and in associational fibers in armadillo, tree shrew, vole, and PWD tissue; FXGs were not detected in associational fibers in opossum. FXGs colocalized with FMRP signal. DAPI (blue) used as a nuclear stain. Arrows indicate FXGs. Scale bar=50 µm, 14 µm in insets.

Our past studies of FXGs in the adult mouse brain focused on C57Bl/6 mice, a strain of the *Mus musculus domesticus* subspecies. To determine whether the absence of FXGs from the adult hippocampus may be a specific feature of *M. musculus domesticus*, we asked whether FXGs were abundant in the hippocampus of adult PWD mice, a strain of the *M. musculus musculus* subspecies that diverged from *M. musculus domesticus* approximately 1 million years ago (Montagutelli et al., 1991; Gregorová and Forejt, 2000). As with the C57Bl/6 mice, we only rarely detected FXGs indicating that FXGs are not common in adult hippocampus of multiple *M. musculus* subspecies.

### FMRP is a conserved component of hippocampal FXGs

The function and regulation of RNA granules are heavily dependent on the RNA binding proteins found within those granules. FXGs in the mouse forebrain contain FMRP and FXR2P, with a region-selective subset also containing FXR1P (Christie et al., 2009; Chyung et al., 2018). We therefore asked whether FMRP and FXR2P are conserved components of the hippocampal FXGs found in other species. We performed immunohistochemistry for FXR2P and FMRP on hippocampal sections. We found that FMRP was a constituent of essentially all FXGs in all of the examined species - tree shrews, armadillos, opossums, voles, and PWD mice-regardless of the relative FXG abundance in each species (Figure 2). FXR2P and FMRP are therefore conserved components of FXGs.

### FXGs persist in mossy fibers in adult voles and Peromyscus

Our survey, along with our past studies of mice and rats (Akins et al., 2017), revealed that mossy fiber FXG abundance varies dramatically between species. To better understand factors that might contribute to this variance, we investigated FXG expression within the Rodentia family which comprises a vast number of species that vary in body size, behavior, and habitat and that together make up approximately 40% of all mammalian species. Mice and rats both express hippocampal FXGs as juveniles but diverge in their adult expression patterns. We therefore examined both juveniles (P21) and young adults (P120) to capture FXG expression at each of the relevant developmental stages. We started with voles and California deer mice (*Peromyscus*) from the Cricetidae family, which diverged from the Muridae family containing mice and rats around 25 million years ago (Steppan et al., 2004). We performed immunohistochemistry on hippocampal tissue sections for FXR2P and identified FXGs using standardized approaches as described in Methods. We quantified FXG abundance in dentate mossy fibers as well as in CA3 associational fibers (summarized in Table 1). FXGs were readily detected in these two regions in both species at P21 (Figure 3). At P120, FXGs were evident in mossy fibers at levels comparable to or higher than those seen at P21, a pattern consistent with the developmental profile seen in rats. In contrast, FXG abundance decreased significantly in associational fibers, indicating that the developmental pattern of FXGs varies between the hippocampal circuits.

**Table 1.**
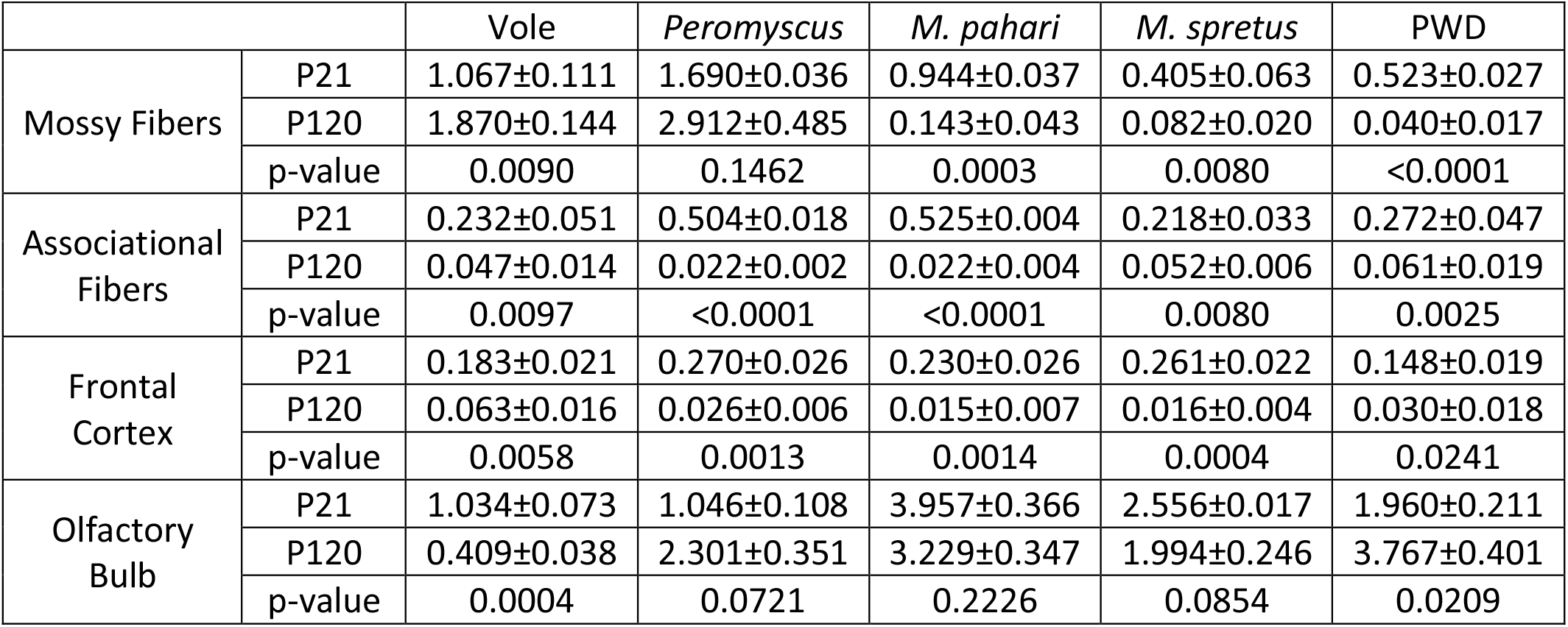
Quantification and analysis of FXG abundance in rodents. FXG abundance across brain regions in voles, *Peromyscus*, *Mus pahari*, *Mus spretus*, and *Mus musculus musculus* (PWD) was quantified and reported as FXG number per 100 µm^2^ (mean±SEM). Statistical analysis by Student’s T-test; n=3 except *Peromyscus* P21 where n=2.

|  |  | Vole | *Peromyscus* | *M. pahari* | *M. spretus* | PWD |
| --- | --- | --- | --- | --- | --- | --- |
| Mossy Fibers | P21 | 1.067±0.111 | 1.690±0.036 | 0.944±0.037 | 0.405±0.063 | 0.523±0.027 |
|  | P120 | 1.870±0.144 | 2.912±0.485 | 0.143±0.043 | 0.082±0.020 | 0.040±0.017 |
|  | p-value | 0.0090 | 0.1462 | 0.0003 | 0.0080 | <0.0001 |
| Associational Fibers | P21 | 0.232±0.051 | 0.504±0.018 | 0.525±0.004 | 0.218±0.033 | 0.272±0.047 |
|  | P120 | 0.047±0.014 | 0.022±0.002 | 0.022±0.004 | 0.052±0.006 | 0.061±0.019 |
|  | p-value | 0.0097 | <0.0001 | <0.0001 | 0.0080 | 0.0025 |
| Frontal Cortex | P21 | 0.183±0.021 | 0.270±0.026 | 0.230±0.026 | 0.261±0.022 | 0.148±0.019 |
|  | P120 | 0.063±0.016 | 0.026±0.006 | 0.015±0.007 | 0.016±0.004 | 0.030±0.018 |
|  | p-value | 0.0058 | 0.0013 | 0.0014 | 0.0004 | 0.0241 |
| Olfactory Bulb | P21 | 1.034±0.073 | 1.046±0.108 | 3.957±0.366 | 2.556±0.017 | 1.960±0.211 |
|  | P120 | 0.409±0.038 | 2.301±0.351 | 3.229±0.347 | 1.994±0.246 | 3.767±0.401 |
|  | p-value | 0.0004 | 0.0721 | 0.2226 | 0.0854 | 0.0209 |

**Figure 3.**
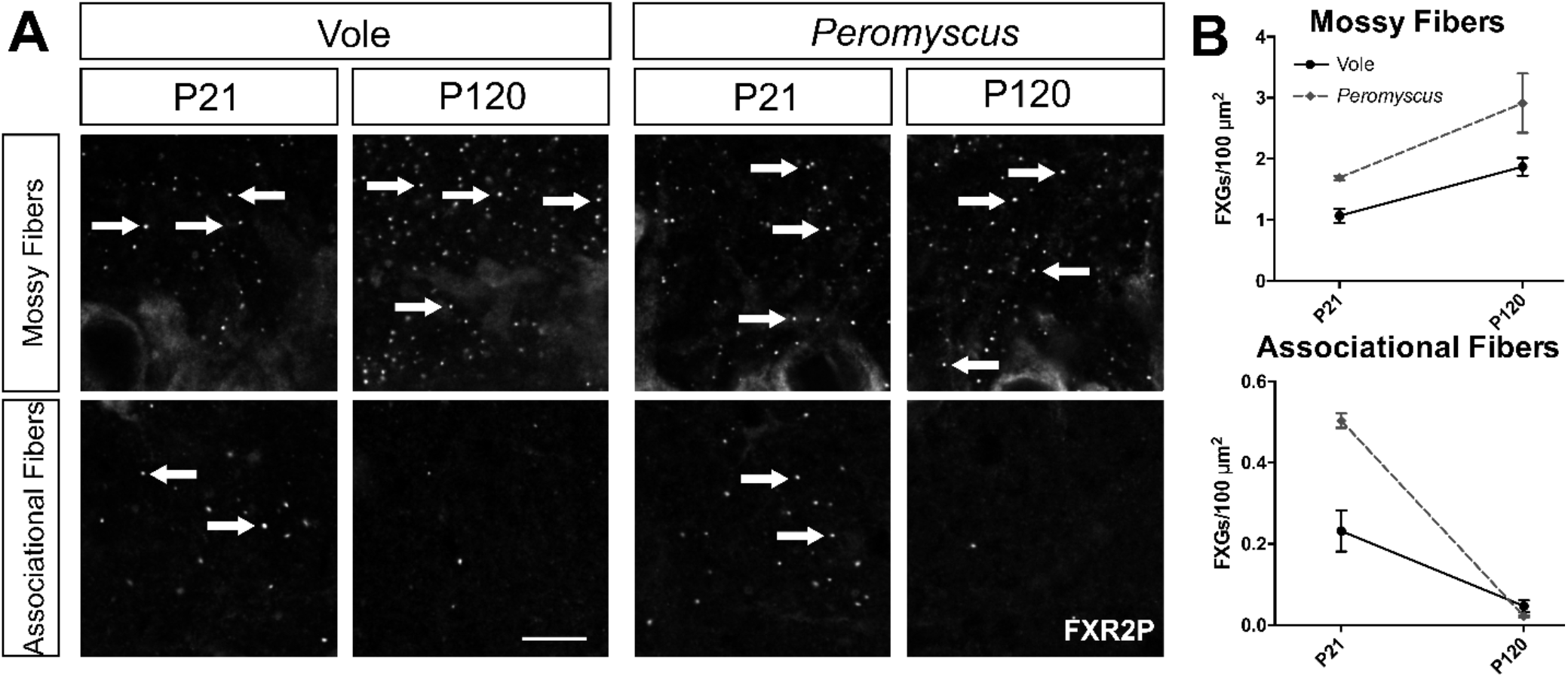
Mossy fiber FXGs persist into adulthood in some members of the Rodentia family. **(A)** Confocal micrographs of immunostaining for FXR2P in hippocampal mossy and associational fibers. FXGs are abundant at P21 in vole and *Peromyscus* in both mossy and associational fibers, but while mossy fiber FXG abundance stays high through P120, it decreases drastically in associational fibers. **(B)** Quantification of FXG abundance in juvenile and adult vole and *Peromyscus* in mossy fibers and associational fibers. See Table 1 for FXG quantification and statistics values; n=3 except *Peromyscus* P21 where n=2. Arrows indicate FXGs. Scale bar=10 µm.

### Hippocampal FXGs are lost in adulthood in the Mus genus

Our findings that adult hippocampal FXGs are rare in two different *M. musculus* subspecies, but abundant in rat, vole, and *Peromyscus* suggested that this difference may represent a species-specific divergence in FXG regulation that reflects differences in how the species live in the wild. However, lab mice have also been subjected to extensive controlled breeding for over a century, raising the possibility that the change in regulation may represent a quirk of selective breeding rather than revealing a difference in their native biology. To investigate these possibilities, we examined the developmental profile of hippocampal FXG expression in *M. musculus* subspecies including 6 additional strains of *M. musculus domesticus* (a mix of inbred and outbred strains; DBA2, BALB/C, 129, FVB, CFW, and CD1), *M. musculus musculus* (PWD), and *M. musculus castaneus*. We also extended our analyses to two related species within the *Mus* genus: *M. pahari* and *M. spretus*, which diverged from *M. musculus* approximately 6 million years ago (Thybert et al., 2018) and 2 million years ago (Galtier et al., 2004; Suzuki et al., 2004), respectively. Strikingly, the developmental pattern of hippocampal FXGs was consistent across these *Mus* species, subspecies, and strains. FXGs were present in both mossy and associational fibers in all the examined juveniles but were rare in both circuits in the adults (Figure 4; Table 1). This pattern was unaffected by breeding history, as the examined *Mus* animals included inbred, outbred, and recently wild-derived species and strains (Table 2). Instead, downregulation of FXG abundance in adult hippocampus was conserved throughout the *Mus* genus.

**Table 2.**
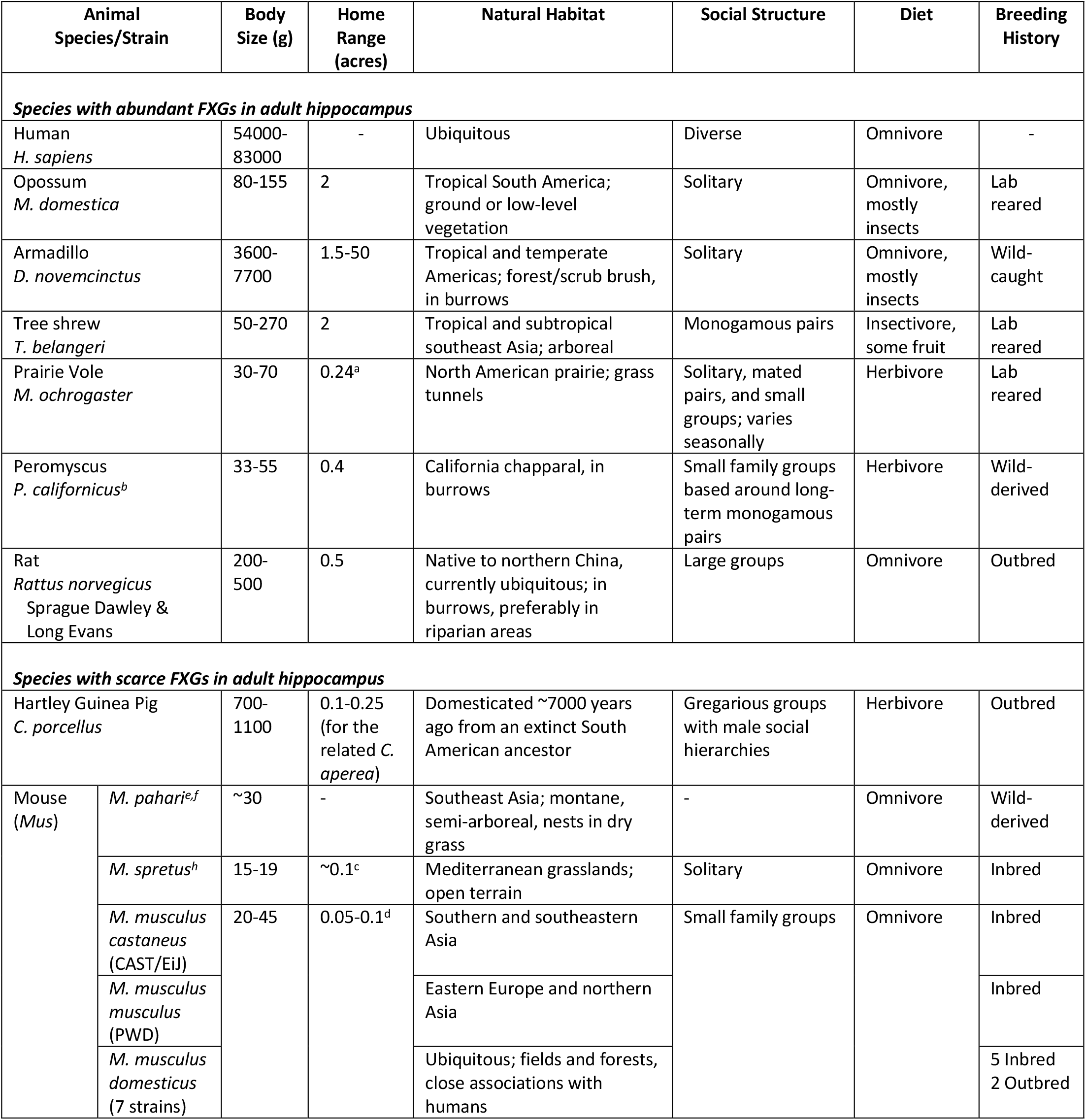
Characteristics of examined species. Information collected from Animal Diversity Web, University of Michigan Museum of Zoology (Myers et al., 2018), except as noted: a: (Linzey and Hammerson, 2015); b: (MacMillen, 1964); c: (Gray et al., 1998); d: (Mikesic and Drickamer, 1992); e: (Molur et al., 2005); f: (Smith and Xie, 2008); g: (Montoto et al., 2011); h: (Palomo et al., 2009)

**Table 3.** Quantification and analysis of FXG abundance in guinea pigs. FXG abundance across brain regions in guinea pigs was quantified and reported as number of FXGs per 100 µm^2^ (mean±SEM). Statistical analysis by ANOVA with multiple comparisons; n=3 except olfactory bulb where n=1 at P2 n=1; n=2 at P21 and P120. There was no statistical difference between P21 and P120 for any region.

|  | **Mossy fibers (CA4)**<br>(ANOVA p=0.0068) | **Mossy fibers (CA3)**<br>(ANOVA p=0.0023) | **Associational fibers**<br>(ANOVA p=0.0237) | **Frontal Cortex**<br>(ANOVA p=0.0070) | **Olfactory Bulb** |
| --- | --- | --- | --- | --- | --- |
| P2 | 0.434±0.032 | 0.276±0.023 | 0.279±0.035 | 0.149±0.001 | 0.450 |
| P21 | 0.252±0.037<br>(p=0.0124 vs P2) | 0.139±0.026<br>(p=0.0073 vs P2) | 0.167±0.010<br>(p=0.0373 vs P2) | 0.092±0.014<br>(p=0.0119 vs P2) | 0.274±0.007 |
| P120 | 0.230±0.022<br>(p=0.0108 vs P2) | 0.098±0.010<br>(p=0.0029 vs P2) | 0.178±0.014<br>(p=0.0402 vs P2) | 0.086±0.009<br>(p=0.0118 vs P2) | 0.319±0.079 |

**Figure 4.**
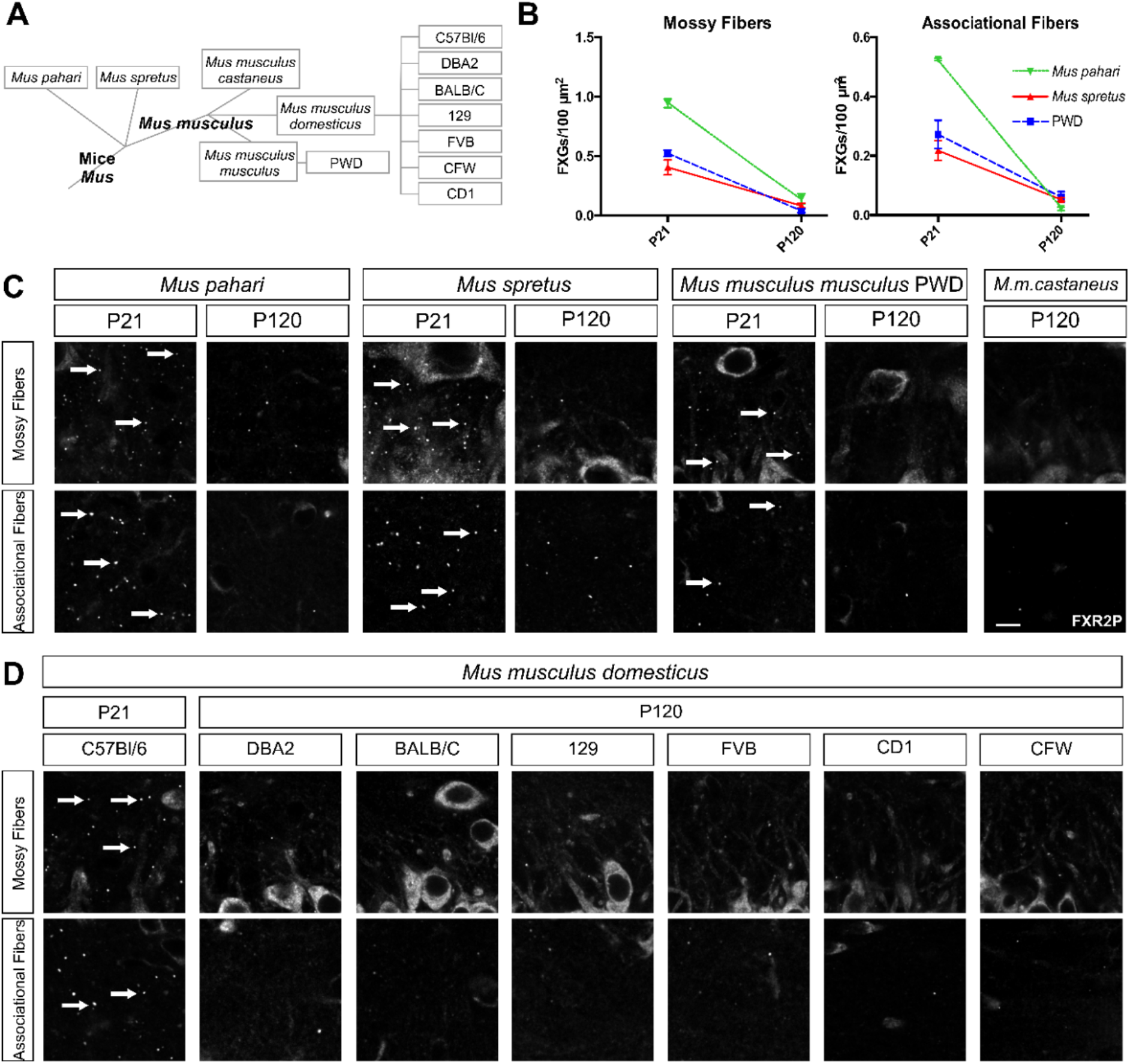
Hippocampal FXGs are lost in adulthood in the *Mus* genus. **(A)** Taxonomic key of the representative members of the Mus genus. **(B)** Quantification of FXG abundance in juveniles and adults in *Mus pahari*, *Mus spretus*, and *Mus musculus musculus* (PWD) in both mossy fibers and associational fibers. While FXGs are abundant at P21 in both regions, they are nearly undetectable by P120 in all *Mus* species examined. See Table 1 for FXG quantification and statistics values. **(C)** Confocal micrographs of immunostaining for FXR2P in hippocampal mossy fibers and associational fibers of *M. pahari*, *M. spretus*, PWD, and *Mus musculus castaneus*. (B) FXGs are abundant in both regions at P21 in *Mus pahari*, *Mus spretus*, and PWD but are nearly absent at P120. FXGs are sparsely detected in *M.m. castaneus* at P120 in both regions. **(D)** Confocal micrographs of immunostaining for FXR2P in hippocampal mossy and associational fibers of 7 strains of *Mus musculus domesticus*, including juvenile C57Bl/6, adult inbred strains DBA2, BALB/C, 129, and FVB, and adult outbred strains CFW and CD1. FXGs are only sparsely detected in both regions in the adult *M.m.domesticus* strains compared to juvenile C57Bl/6. Arrows indicate FXGs. n=3. Scale bar=10 µm.

### Frontal cortex and olfactory bulb FXG developmental patterns are consistent across species

Previous findings have indicated that, in contrast to the hippocampus, the FXG developmental profile is similar between mice and rats in frontal cortex (where both species exhibit a developmental decrease in FXG abundance) and olfactory bulb (where FXGs are expressed across the lifespan) (Akins et al., 2017). To determine if these observed patterns are consistent across species, we quantified FXG abundance in frontal cortex and olfactory bulb in vole, *Peromyscus*, *M. pahari*, *M. spretus*, and PWD mice at P21 and P120 (Figure 5). FXGs were found in these circuits in all species examined. Consistent with the protein composition seen in past studies in mice, essentially all granules in both brain regions contained both FXR2P and FMRP. In all species FXG abundance declined significantly in the frontal cortex between P21 and P120.

**Figure 5.**
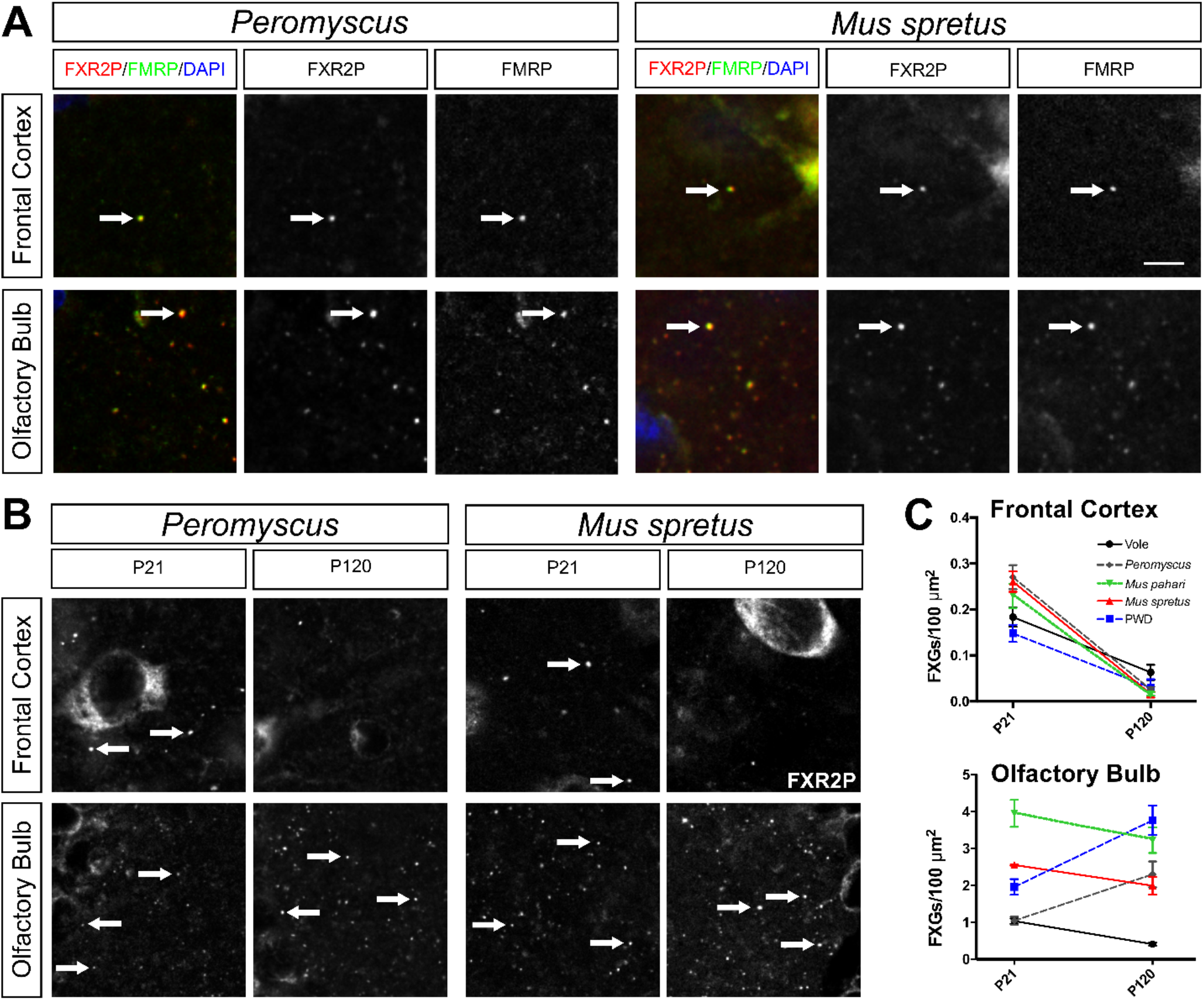
Frontal cortex and olfactory bulb FXG developmental pattern throughout the Rodentia family. **(A)** Confocal micrographs of coimmunostaining of FXGs, indicated by FXR2P (red) with FMRP (green) in frontal cortex and olfactory bulb of P120 *Peromyscus* and *Mus spretus*. FMRP colocalized with FXGs in both regions at P21 and P120 in all species examined (vole, PWD, *Peromyscus*, *Mus pahari*, and *Mus spretus*, data not shown). DAPI (blue) used as a nuclear stain. **(B)** Confocal micrographs of immunostaining for FXR2P in frontal cortex and olfactory bulb of *Peromyscus* and *Mus spretus* at P21 and P120. Arrows indicate FXGs. **(C)** Quantification of frontal cortex and olfactory bulb FXG abundance in *Peromyscus*, vole, *Mus pahari*, *M. spretus*, and PWD at P21 and P120. In frontal cortex of all species examined, FXG abundance declines drastically by P120 compared to P21. In olfactory bulb, FXG abundance did not change with age in vole, *M. pahari*, or *M. spretus*, but were higher in adults than juveniles in *Peromyscus* and PWD. See Table 1 for FXG quantification and statistics values. n=3 except *Peromyscus* P21 where n=2. Arrows indicate FXGs. Scale bar=6 µm (A), 10 µm (B).

**Figure 6.**
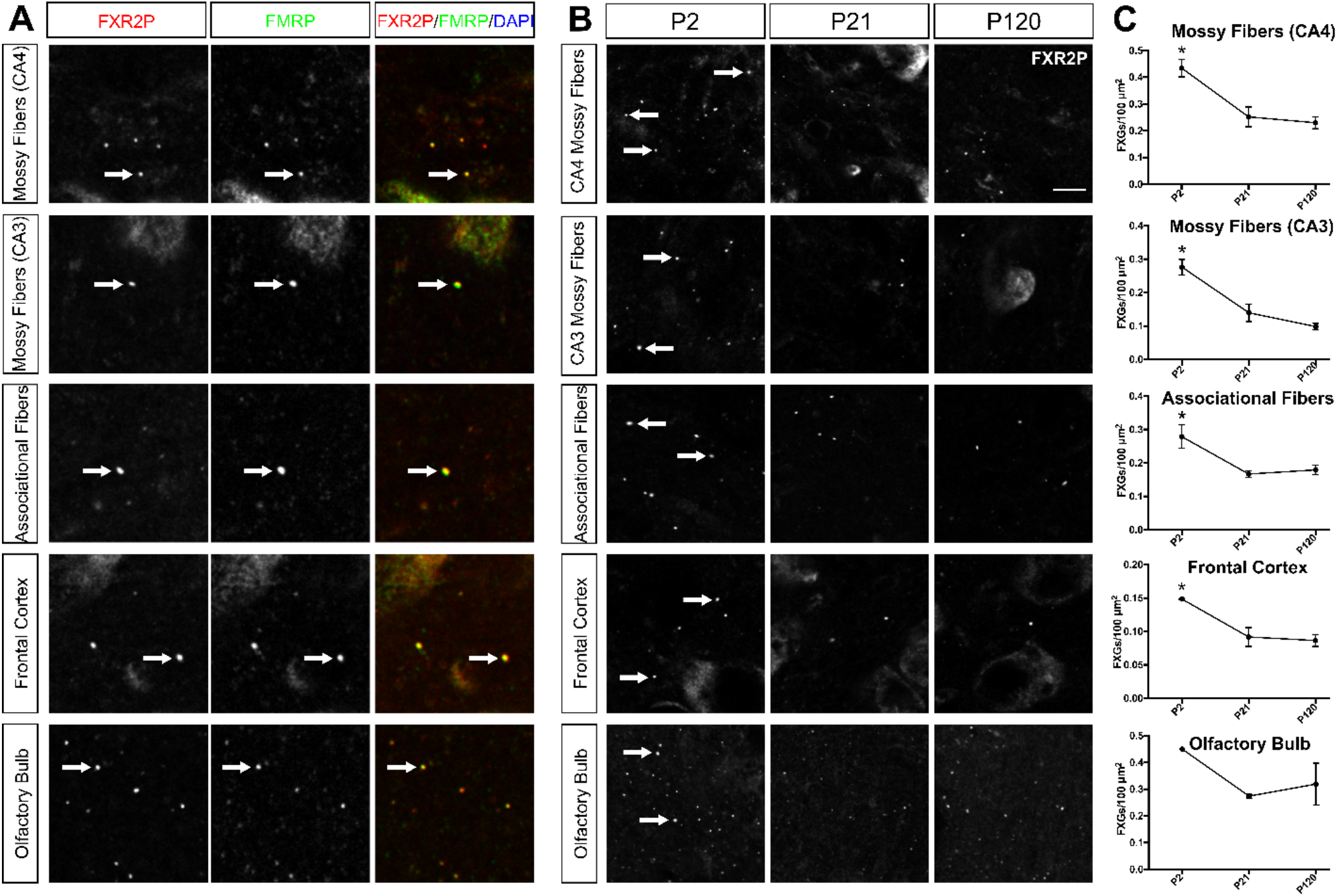
FMRP-containing FXGs are abundant across the brain in neonatal guinea pigs and decline with age. **(A)** Confocal micrographs of coimmunostaining for FXGs, indicated by FXR2P (red) and FMRP (green) in mossy fibers, associational fibers, frontal cortex, and olfactory bulb of P120 guinea pig. FMRP colocalized with FXGs in all four regions at P120 as well as P2 and P21 (data not shown). DAPI (blue) used as a nuclear stain. Arrows indicate FXGs. **(B)** Confocal micrographs of immunostaining for FXGs, indicated by FXR2P, in mossy fibers in the CA3 and CA4 regions, associational fibers, frontal cortex, and olfactory bulb in guinea pig at P2, P21, and P120. **(C)** Quantification of FXG abundance. FXG abundance decreased significantly in all hippocampal regions and frontal cortex between the neonate (P2) and juvenile (P21), then remained stable into adulthood (P120). See Table 3 for FXG quantification and statistics values. n=3 for all regions except olfactory bulb where P2 n=1, P21 and P120 n=2. Scale bar=6 µm (A), 10 µm (B).

Although FXG abundance in the olfactory bulbs of all species remained high at both ages, we did observe small but significant changes in most species. FXG abundance remains steady in *Peromyscus*, *M. pahari*, and *M. spretus* but decreased with age in voles and increased with age in PWD mice (Figure 5B-C, Table 1). Both rats and C57Bl/6 mice exhibit a developmental increase in olfactory bulb FXG abundance up to a peak that is then followed by a decrease to the levels seen in adults (Christie et al., 2009; Akins et al., 2017). The developmental timing of these peaks differs – for rats it occurs at around P60 while for C57Bl/6 mice it occurs at about P30. The species-dependent differences in FXG abundance seen in the present study may reflect similar variability in the timing of peak FXG abundance. Taken together, our findings from hippocampus, frontal cortex, and olfactory bulb indicate that the developmental profile of FXG expression in forebrain circuits is largely conserved across species with the single exception of dentate mossy fibers.

### Species with small home range sizes lack mossy fiber FXGs in adulthood

The divergence in adult hippocampal FXG abundance in the *Mus* lineage raises the possibility that these animals were subjected to different environmental constraints that led to a diminished requirement for FXG function in the adult hippocampus. We examined potentially relevant factors including body size, home range size, natural habitat, social structure, diet, breeding history, and housing conditions (Table 2). Among these, *Mus* were distinguished only by their home range size, which is substantially smaller for *Mus* than for the other examined animals.

To investigate the potential contribution of home range size, we investigated FXG expression in guinea pigs, who also occupy small home ranges. However, in contrast to the other, altricial rodent species examined here, guinea pigs are precocial. Our past studies in mice revealed that FXGs are rare until approximately P15, coinciding with eye opening and increased exploration (Christie et al., 2009). Because guinea pigs can fully explore their environment essentially immediately after birth, we hypothesized that they might express FXGs earlier than mice. We therefore investigated FXG expression in P2 neonates in addition to P21 juveniles and P120 adults. We observed FXGs in P2 animals in axons of all examined brain regions including mossy fibers, CA3 associational fibers, olfactory sensory neuron axons in olfactory bulb, and axons innervating frontal cortex. Because guinea pig hippocampus is more elaborated than that in other rodents with defined CA4 and CA3 regions, we quantified FXGs in dentate mossy fibers in both of these regions. In all five regions, abundance declined significantly from P2 to P21, attaining levels consistent with those seen at P120. Further, as in all examined species, FXR2P and FMRP were components of FXGs in all of these brain regions. The decrease in mossy fiber FXGs in guinea pigs differs from the developmental increase seen in other species and instead echoes, although less dramatically, the developmental decrease seen in *Mus* species.

## Discussion

In this study, we investigated FXG expression across mammalian species as a means to address whether these granules are likely to play fundamental, conserved roles. FXGs were found in the brains of every examined species, including both placental and nonplacental mammals. Many aspects of these granules were conserved including the RNA binding proteins they comprise, the circuits in which they are found, and the developmental timing of FXG expression in olfactory bulb and neocortex. Interestingly, one aspect that diverged among species was the developmental window in which FXGs were found in hippocampus, with FXGs found in the adult hippocampus of most but not all mammals. Investigation of this divergence within rodents suggested that the species that lose FXG expression in the adult hippocampus may be those that occupy small home ranges in the wild. Together, these findings indicate that FXGs are broadly conserved in the mammalian brain and that FXG-regulated translation likely plays important roles in controlling the neuronal proteome.

Local protein synthesis following transport of mRNAs to particular subcellular locations is a fundamental mechanism for controlling the spatial distribution of proteins within cells that occurs not only in eukaryotes but also in bacteria (Martin and Ephrussi, 2009; Nevo-Dinur et al., 2011; Vazquez-Pianzola and Suter, 2012; Jung et al., 2014; Buxbaum et al., 2015). The RNPs and their constituent RNA binding proteins that are associated with the transport and translational control of localized mRNAs are specific to particular lineages with, for example, some found only in yeast or in animals (Claußen and Suter, 2005; Heym and Niessing, 2012; Vazquez-Pianzola and Suter, 2012). Within animals, the FXR protein family is widely conserved and consistently controls the translation of localized mRNAs within neurons (Bassell and Warren, 2008). Among the FXR protein-containing complexes, FXGs are unique with regard to their neuron type specificity and subcellular distribution solely within axons. The findings presented here reveal that this specificity is consistent across a remarkably diverse collection of mammalian species indicating that the mechanisms that govern FXG formation and localization are highly conserved. Moreover, the widespread conservation of these axonal RNPs suggests that local axonal protein synthesis, a phenomenon that has only recently been widely studied, is an ancient, conserved mechanism for regulating the axonal proteome in the mammalian brain.

Several lines of evidence suggest that FXG-regulated translation modulates experience-dependent synaptogenesis. FXGs are most abundant in axons that are undergoing experience-dependent synapse formation and elimination. For example, in adult animals FXGs are consistently seen across all species in olfactory sensory neuron axons in the olfactory bulb (Christie et al., 2009; Akins et al., 2017). These neurons are generated throughout life and exhibit a remarkable degree of experience-dependent structural plasticity during initial circuit integration that continues long after the axons have formed synapses and reached maturity (Cheetham et al., 2016). Lesion studies of the olfactory neurons showed that FXGs are most abundant as newly formed neurons are forming synapses (Christie et al., 2009). Interestingly, the developmental onset of FXG expression further points toward a correlation with circuit refinement. In mice and rats, FXGs are rare or absent at birth and don’t appear until approximately two weeks of age (Christie et al., 2009). This timing corresponds to a period of intense sensory exploration of the world since it occurs during the time when visual and auditory information first impinge on the CNS, leading to experience-dependent circuit refinement (Hensch, 2004, 2005; Barkat et al., 2011). In contrast to rats and mice, guinea pigs are precocial and therefore receive and process sensory information and eat solid food shortly after birth (Künkele and Trillmich, 1997). Notably, FXGs were found in brains from neonatal guinea pigs, further supporting a correlation between FXG onset and initial sensory experience. Together, these findings suggest that FXGs play conserved roles in circuits during periods of robust experience-dependent synaptogenesis.

In addition to first appearing as circuits are forming, FXGs are developmentally downregulated in most brain regions, including cerebellum and neocortex, after robust experience-dependent circuit refinement has concluded (Christie et al., 2009; Akins et al., 2017). In contrast, FXGs persist into adulthood in both olfactory bulb and hippocampus in most species. While lesion studies showed a link between neurogenesis and FXG expression in the olfactory bulb (Christie et al., 2009), FXG expression in the hippocampus is independent of neurogenesis (Akins et al., 2017), suggesting an ongoing role for FXG-regulated translation in mature axons in this circuit. Interestingly, adult FXG expression was lost in the *Mus* lineage sometime after it diverged from the *Rattus* lineage, even though expression in the juveniles is similar between mice and rats. This deviation could merely reflect random genetic variation, but it also raises the possibility that the organismal roles that hippocampal FXGs support are less prominent in adult mice than in adults of other mammalian species. A survey of factors that may have led to this deviation pointed towards the comparatively small home range occupied by mice in the wild, suggesting that ecological factors may contribute to whether FXGs persist or are absent from adult hippocampus of particular species. Consistent with this idea, guinea pigs, which also occupy very small home ranges in the wild, were the only examined species outside the *Mus* genus that exhibited a developmental decline in FXG expression in hippocampal mossy fibers. As discussed above, the developmental timing of FXGs, as well as their mRNA cargoes, suggest that FXGs may modulate the formation and modification of synapses during circuit wiring (Christie et al., 2009; Akins et al., 2017; Chyung et al., 2018). Sustained expression of FXGs in the adult hippocampus may support the continuous circuit rewiring that occurs as animals navigate a large and changing world. The relative paucity of these structures in adult mice and guinea pigs may reflect the smaller territories inhabited by individuals of these species.

Axonal FMRP functions are varied and include regulation of ion channel function and localization as well as regulation of protein synthesis during axon outgrowth and synapse formation (Li et al., 2009; Deng et al., 2013; Ferron et al., 2014; Zimmer et al., 2017). Loss of these functions likely contributes to the cognitive and neurological symptoms seen in FXS patients. In particular, the presence of FXGs, and their associated translational machinery, in the adult human hippocampus points towards a role for FMRP-regulated local protein synthesis in this circuit in humans. The majority of our mechanistic understanding of FMRP and how its loss impacts synaptic function come from the study of mouse models of Fragile X. In terms of presynaptic FMRP in mature hippocampal circuits, however, mice are not representative of the vast majority of mammalian species including humans. Elucidating how FXG-associated presynaptic FMRP contributes to adult circuit function, and how its loss may impact adult Fragile X patients, therefore requires studies in species outside the *Mus* genus. For example, *Peromyscus* and rats both engage in complex social behaviors and provide ideal models for investigating the relationship between such behaviors and the regulation and function of FXGs. Moreover, *Fmr1* null rats (Hamilton et al., 2014) provide a basis for understanding FXG functions in adult mammals in general as well as a more clinically relevant model than *Fmr1* null mice for the investigation of FXG dysregulation in human FXS patients.

## Methods

### Animals

All work with animals was performed in accordance with protocols approved by the Institutional Animal Care and Use Committee of Drexel University and the respective institutions that donated tissue. Male and female animals did not exhibit differences in FXG expression or composition at any age; therefore, all studies used a mix of males and females. We collected tissue from the following animals: *Mus spretus* (SPRET/EiJ), *Mus pahari*, *Mus musculus castaneus* (CAST/EiJ), *Mus musculus musculus* (PWD) (all from Jackson Laboratory, Bar Harbor, ME), and *Mus musculus domesticus* strains (DBA2, BALB/C, 129, FVB, CFW, CD1; Charles River Laboratories, Wilmington, MA). Animals were deeply anaesthetized by an intraperitoneal injection of ketamine/xylazine/acepromazine before intracardiac perfusion with room temperature HBS (0.1M HEPES, pH 7.4; 150 mM sodium chloride) containing 1U/mL heparin and 0.5% sodium nitrite followed by perfusion with room temperature PBS (0.1M phosphate, pH 7.4; 150 mM sodium chloride) containing 4% paraformaldehyde. After perfusion, intact brains were carefully removed and postfixed overnight in the perfusate. After washing in PBS, brains were transferred to PBS containing 30% sucrose until the brains sank for cryoprotection. Brains were then embedded in OCT medium by rapid freezing and stored at −80°C until sectioned. Free-floating coronal or sagittal sections of OCT-embedded brains were prepared using a Leica cryostat at 40 μm and either used the same day or stored at 4°C in PBS containing 0.02% sodium azide.

Perfused brains were generously donated by Dr. Elizabeth Becker (St. Joseph’s University; *Peromyscus*), Dr. Bruce Cushing (The University of Texas at El Paso; vole), Dr. Jeffery Padburg (University of Central Arkansas; armadillo), and Dr. Leah Krubitzer (University of California, Davis; opossum and tree shrew). Perfused guinea pig brains were purchased from Hilltop Lab Animals (Scottdale, PA). Donated and purchased tissue was perfused with 4% paraformaldehyde prior to being shipped and stored at 4 °C in PBS containing 0.02% sodium azide and processed as above starting with cryoprotection in PBS with 30% sucrose.

### Immunohistochemistry

Tissue was first washed three times in PBS (10 mM phosphate pH 7.4; 150 mM NaCl). To improve antibody accessibility to epitopes in tissue, sections were then heated in 0.01M sodium citrate (pH 6.0) for 10 min at 99°C. Tissue was then treated with blocking solution [PBST (10 mM phosphate buffer, pH 7.4, and 0.3% Triton X-100) and 1% blocking reagent (Roche)] for 30min to occupy nonspecific binding sites. Sections were then incubated with blocking solution plus primary antibody overnight before being washed for 5 min with PBST. For secondary detection, tissue was incubated with appropriate labelled antibodies in blocking solution for 1 h. Tissue was then washed for 5 min with PBST, mounted in 4% n-propylgallate, 85% glycerol, 10 mM phosphate pH 7.4, then coverslipped and sealed with nail polish. Sections were imaged using a Leica SPE confocal microscope.

### Antibody Characterization

*Antibody Characterization:*
| **Primary Antibody** | **Source** | **Target** | **RRID** | **Primary Dilution** | **Secondary antibody** |
| --- | --- | --- | --- | --- | --- |
| Mouse anti-FXR2P | Monoclonal A42 from hybridoma supernatant | Raised against human FXR2P amino acids 12-426; recognizes a single band of the appropriate size by Western and does not cross react with either FMRP or FXR1P (Zhang et al., 1995) | AB_2737297 | 1:300 | Alexa-conjugated goat anti-mouse IgG1 (Life Technologies; 1:1000) |
| Mouse anti-FMRP | Monoclonal 2F5-1 from concentrated hybridoma supernatant | Raised against amino acids 1–204 of human FMRP. Recognizes bands of the appropriate sizes by Western, does not give signal in either blots or sections derived from Fmr1 null mice (Gabel et al., 2004; Christie et al., 2009) | AB_10805421 | 5 µg/ml | Alexa-conjugated goat anti-mouse IgG2b (Life Technologies; 1:1000) |

### FXG Composition

Fluorescent micrographs of brain sections immunostained for FXR2P and FMRP were collected as above. FXGs were identified based on morphology using FXR2P signal as previously described (Christie et al., 2009; Akins et al., 2012, 2017; Chyung et al., 2018). Each FXR2P-containing FXG was manually annotated as to whether it colocalized with FMRP signal.

### FXG Quantification

Epifluorescent micrographs of FXR2P immunostaining were captured from three to five sections of each condition (age, species, and brain region as indicated in Results) with a Hamamatsu Orca-R^2^ camera coupled to a Leica DMI4000 B microscope using a 40x oil objective (NA=1.15). These images were processed using the FIJI build of ImageJ with a rolling ball background subtraction with a radius of 7 pixels followed by a median filter with a radius of 2 pixels. All pixels less than 1.5 standard deviations above the image background intensity were removed from further processing. Auto threshold was set using maximum entropy of the remaining pixels for FXG identification. Granules were counted manually using previously described criteria (Christie et al., 2009; Akins et al., 2012) with pixels inappropriately identified as FXGs (e.g., autofluorescent blood vessels) manually discarded from counts. Graphs and statistical analyses were produced using Prism 6.0 (Graphpad). Quantifications are expressed as mean #x00B1; SEM.

### Figure Preparation

Images used in figures were adjusted for brightness, contrast, and gamma using Photoshop CS6 13.0.6 (Adobe). Figures were prepared using Corel Draw X7 (Corel).

## Acknowledgements

Many thanks to colleagues who donated animal tissue: Dr. Elizabeth Becker (St. Joseph’s University; *Peromyscus*), Dr. Bruce Cushing (The University of Texas at El Paso; vole), Dr. Jeffery Padburg (University of Central Arkansas; armadillo), and Dr. Leah Krubitzer (University of California, Davis; opossum and tree shrew). Thanks to Dr. Dylan Cooke for helping arrange tissue donations, and to Duy Le, Sachi Desai, Anshul Ramanathan, and Eric Gebski for technical assistance.

## References

Akins MR, Berk-Rauch HE, Kwan KY, Mitchell ME, Shepard KA, Korsak LIT, Stackpole EE, Warner-Schmidt JL, Sestan N, Cameron HA, Fallon JR (2017) Axonal ribosomes and mRNAs associate with fragile X granules in adult rodent and human brains. Hum Mol Genet 26:192–209.

Akins MR, LeBlanc HF, Stackpole EE, Chyung E, Fallon JR (2012) Systematic mapping of fragile X granules in the mouse brain reveals a potential role for presynaptic FMRP in sensorimotor functions. J Comp Neurol 520:3687–3706.

Barkat TR, Polley DB, Hensch TK (2011) A critical period for auditory thalamocortical connectivity. Nat Neurosci 14:1189–1194.

Bassell GJ, Warren ST (2008) Fragile X Syndrome: Loss of Local mRNA Regulation Alters Synaptic Development and Function. Neuron 60:201–214.

Buxbaum AR, Haimovich G, Singer RH (2015) In the right place at the right time: visualizing and understanding mRNA localization. Nat Rev Mol Cell Biol 16:95–109.

Cheetham CEJ, Park U, Belluscio L (2016) Rapid and continuous activity-dependent plasticity of olfactory sensory input. Nat Commun 7:10729.

Christie SB, Akins MR, Schwob JE, Fallon JR (2009) The FXG: A Presynaptic Fragile X Granule Expressed in a Subset of Developing Brain Circuits. J Neurosci 29:1514–1524.

Chyung E, LeBlanc HF, Fallon JR, Akins MR (2018) Fragile X granules are a family of axonal ribonucleoprotein particles with circuit-dependent protein composition and mRNA cargos. J Comp Neurol 526:96–108.

Claußen M, Suter B (2005) BicD-dependent localization processes: from Drosophilia development to human cell biology. Ann Anat - Anat Anz 187:539–553.

Darnell JC, Klann E (2013) The translation of translational control by FMRP: therapeutic targets for FXS. Nat Neurosci 16:1530–1536.

Deng P-Y, Rotman Z, Blundon JA, Cho Y, Cui J, Cavalli V, Zakharenko SS, Klyachko VA (2013) FMRP Regulates Neurotransmitter Release and Synaptic Information Transmission by Modulating Action Potential Duration via BK Channels. Neuron 77:696–711.

Ferron L, Nieto-Rostro M, Cassidy JS, Dolphin AC (2014) Fragile X mental retardation protein controls synaptic vesicle exocytosis by modulating N-type calcium channel density. Nat Commun 5 Available at: http://www.nature.com/ncomms/2014/140407/ncomms4628/full/ncomms4628.html [Accessed May 6, 2014].

Gabel LA, Won S, Kawai H, McKinney M, Tartakoff AM, Fallon JR (2004) Visual Experience Regulates Transient Expression and Dendritic Localization of Fragile X Mental Retardation Protein. J Neurosci 24:10579–10583.

Galtier N, Bonhomme F, Moulia C, Belkhir K, Caminade P, Desmarais E, Duquesne JJ, Orth A, Dod B, Boursot P (2004) Mouse biodiversity in the genomic era. Cytogenet Genome Res 105:385–394.

Gray SJ, Hurst JL, Stidworthy R, Smith J, Preston R, MacDougall R (1998) Microhabitat and spatial dispersion of the grassland mouse (Mus spretus Lataste). J Zool 246:299–308.

Gregorová S, Forejt J (2000) PWD/Ph and PWK/Ph inbred mouse strains of Mus m. musculus subspecies–a valuable resource of phenotypic variations and genomic polymorphisms. Folia Biol (Praha) 46:31–41.

Hamilton SM, Green JR, Veeraragavan S, Yuva L, McCoy A, Wu Y, Warren J, Little L, Ji D, Cui X, Weinstein E, Paylor R (2014) Fmr1 and Nlgn3 knockout rats: novel tools for investigating autism spectrum disorders. Behav Neurosci 128:103–109.

Hensch TK (2004) Critical period regulation. Annu Rev Neurosci 27:549–579.

Hensch TK (2005) Critical period plasticity in local cortical circuits. Nat Rev Neurosci 6:877–888.

Heym RG, Niessing D (2012) Principles of mRNA transport in yeast. Cell Mol Life Sci 69:1843–1853.

Jung H, Gkogkas CG, Sonenberg N, Holt CE (2014) Remote Control of Gene Function by Local Translation. Cell 157:26–40.

Künkele J, Trillmich F (1997) Are Precocial Young Cheaper? Lactation Energetics in the Guinea Pig. Physiol Zool 70:589–596.

Li C, Bassell G j, Sasaki Y (2009) Fragile X mental retardation protein is involved in protein synthesis-dependent collapse of growth cones induced by Semaphorin-3A. Front Neural Circuits 3:11.

Linzey AV, Hammerson G (2015) Microtus ochrogaster, Prairie Vole. International Union for Conservation of Nature and Natural Resources. Available at: www.iucnredlist.org.

MacMillen RE (1964) Population ecology, water relations, and social behavior of a Southern California semi-desert rodent fauna. University of California Publications Zoology.

Martin KC, Ephrussi A (2009) mRNA Localization: Gene Expression in the Spatial Dimension. Cell 136:719–730.

Mikesic DG, Drickamer LC (1992) Factors Affecting Home-Range Size in House Mice (Mus musculus domesticus) Living in Outdoor Enclosures. Am Midl Nat 127:31–40.

Molur S, Srinivasulu C, Srinivasulu B, Walker S, Nameer PO, Ravikumar L (2005) Status of nonvolant small mammals: Conservation Assessment & Management Plan (C.A.M.P.) Workshop Report. Coimbatore, India: Zoo Outreach Organisation/CBSG-South Asia.

Montagutelli X, Serikawa T, Guénet JL (1991) PCR-analyzed microsatellites: data concerning laboratory and wild-derived mouse inbred strains. Mamm Genome Off J Int Mamm Genome Soc 1:255–259.

Montoto LG, Magaña C, Tourmente M, Martín-Coello J, Crespo C, Luque-Larena JJ, Gomendio M, Roldan ERS (2011) Sperm Competition, Sperm Numbers and Sperm Quality in Muroid Rodents. PLoS One San Franc 6:e18173.

Myers P, Espinosa R, Parr CS, Jones T, Hammond GS, Dewey TA (2018) The Animal Diversity Web (online). Anim Divers Web Available at: https://animaldiversity.org/ [Accessed November 5, 2018].

Nevo-Dinur K, Nussbaum-Shochat A, Ben-Yehuda S, Amster-Choder O (2011) Translation-Independent Localization of mRNA in E. coli. Science 331:1081–1084.

Palomo LJ, Justo ER, Vargas JM (2009) Mus spretus (Rodentia: Muridae). Mamm Species:1–10.

Smith AT, Xie Y (2008) A Guide to the Mammals of China. Princeton University Press. Available at: https://press.princeton.edu/titles/8631.html.

Steppan S, Adkins R, Anderson J (2004) Phylogeny and divergence-date estimates of rapid radiations in muroid rodents based on multiple nuclear genes. Syst Biol 53:533–553.

Suzuki H, Shimada T, Terashima M, Tsuchiya K, Aplin K (2004) Temporal, spatial, and ecological modes of evolution of Eurasian Mus based on mitochondrial and nuclear gene sequences. Mol Phylogenet Evol 33:626–646.

Thybert D et al. (2018) Repeat associated mechanisms of genome evolution and function revealed by the Mus caroli and Mus pahari genomes. Genome Res 28:448–459.

Vazquez-Pianzola P, Suter B (2012) Conservation of the RNA Transport Machineries and Their Coupling to Translation Control across Eukaryotes. Int J Genomics Available at: https://www.hindawi.com/journals/ijg/2012/287852/ [Accessed September 4, 2018].

Zhang Y, O’Connor JP, Siomi MC, Srinivasan S, Dutra A, Nussbaum RL, Dreyfuss G (1995) The fragile X mental retardation syndrome protein interacts with novel homologs FXR1 and FXR2. EMBO J 14.

Zimmer SE, Doll SG, Garcia ADR, Akins MR (2017) Splice form-dependent regulation of axonal arbor complexity by FMRP. Dev Neurobiol 77:738–752.

